# High rate of mutational events in SARS-CoV-2 genomes across Brazilian geographical regions, February 2020 to June 2021

**DOI:** 10.1101/2021.07.10.451922

**Authors:** U.J.B. Souza, R.N. Santos, F.S. Campos, K.L. Lourenço, F.G. Fonseca, F.R. Spilki, Corona-ômica.BR/MCTI Network

**Author notes:** Both authors contributed equally (U.J.B. Souza, R.N. Santos). Correspondence author (F.R.S).

## Abstract

Brazil has been considered as one of the emerging epicenters of the coronavirus pandemic in 2021, experiencing over 3,000 daily deaths caused by the virus at the peak of the second wave. In total, the country has more than 19.2 million confirmed cases of Covid-19, including over 533,509 fatalities. A set of emerging variants arose in the country, some of them posing new challenges for COVID-19 control. The goal of this study was to describe mutational events across samples from Brazilian SARS-CoV-2 sequences openly publicly obtainable on the Global Initiative on Sharing Avian Influenza Data-EpiCoV (GISAID-EpiCoV) platform and generate an index of new mutant by each genome. A total of 16,953 SARS-CoV-2 genomes were obtained and are not proportionally representative of the five Brazilian geographical regions. A comparative sequence analysis was conducted to identify common mutations located at 42 positions of the genome (38 were in coding regions whereas two in 5’ and two in 3’ UTR). Moreover, 11 were synonymous and 27 missense variants, and more than 44.4% were located in the spike gene. Across the total of single nucleotide variations (SNVs) identified, 32 were found in genomes obtained from all five Brazilian regions. While a high genomic diversity is reported in Europe given the large number of sequenced genomes, Africa is emerging as a hotspot for new variants. In South America, Brazil and Chile, rates are similar to those found in South Africa and India, giving enough space to generate new viral mutations. Genomic surveillance is the central key to identifying the emerging variants of SARS-CoV-2 in Brazil and has shown that the country is one of the “hotspots” in the generation of new variants.

## 1) Introduction

In December of 2019, at Wuhan, China, a novel betacoronavirus was first detected. Coronavirus Disease 2019 (COVID-19) (Andersen et al., 2020) has developed into a global pandemic, causing waves of epidemics, infecting over 185 million people and 4 million deaths globally by July 10, 2021 (WHO, 2021). The local profile outbreaks were shaped by measures of restrictions, including lockdown, commerce limitations and travel control. The viral spreading has led scientists to investigate genomic epidemiology, which plays a central role in characterizing and understanding the emergence of viruses (Deng et al., 2020; Holmes and Rambaut, 2004; Plowright et al., 2017). The SARS-CoV-2 single stranded RNA is 29.9 kb in size and has positive coding orientation, encoding four major structural proteins on its 3’ end: spike (S), envelope (E), membrane (M) and nucleocapsid (N). These proteins are essential to virus entry into cells and virus particle formation (Bakhshandeh et al., 2021; Souza et al., 2021).

NGS-based SARS-CoV-2 genome characterizations have revolutionized the scale and depth of variant analysis worldwide. Never before has a viral genome been sequenced so globally in such a short time. Even so, numerous reports revealed potential adaptations at the nucleotide (nt), amino acid (aa) and structural heterogeneity of viral proteins, particularly in the S protein (Sardar et al., 2020). At the moment, the world’s concern is focused in about four functionally well-defined variants, B.1.1.7 (Alpha), B.1.351 (Beta), P.1 (Gamma) and B.1.617.2 (Delta) associated with viral fitness changes (Faria et al., 2021; Hodcroft, 2021; McCallum et al., 2021; Tang et al., 2021; Tegally et al., 2021).

The mutation rate in SARS-CoV-2 is about 10^4 replacement of base pairs per year, and possible variations may appear in each replication cycle. In the context of investigating evolutionary events it is possible to compare single-nucleotide polymorphisms (SNPs) in RNA sequences because mutations in coronaviruses occur from RdRp mistakes during viral genome replication (Bakhshandeh et al., 2021; Domingo et al., 1997; Ma et al., 2015).

The spike protein of SARS-CoV-2 contains an N-terminal S1 subunit and a C-terminal membrane proximal to the S2 subunit. The N-terminal domain (NTD) located in the portion S1A, recognizes carbohydrates, such as sialic acid and it is responsible for the attachment of the virus to the host cell surface. The receptor-binding domain (RBD) in the S1B portion interacts with the human ACE-2 receptor. Between S1 and S2 there is a PRRA sequence motif that functions as a furin cleavage site. At S2 the virus has a second cleavage site which also participates in the viral entry into host cells (Guruprasad et al., 2021). The mutation D614G in S, for instance, is a frequently identified mutation that has been associated with increased virus transmissibility and infectivity, and is possibly one of the origins of the widely prevalent B1.1 branch in many countries. Nonetheless, the mutation does not seem to alter the antigenicity of the S protein (Yurkovetskiy et al., 2020) and was not associated with any changes in disease severity (Korber et al., 2020; Plante et al., 2020).

In this study, we evaluated the mutational events across samples from publicly available SARS-CoV-2 sequences available in GISAID in the last year (from February 2020 to June 2021) of pandemics in the country. Moreover, we propose a mathematical relation between new mutants *versus* sequenced genomes. This analysis is fundamental to understand the changes in the viral genome leading to alterations in viral fitness and transmissibility across the population.

## 2) Material and Methods

### 2.1) Data Retrieval

Whole genome sequences of SARS-CoV-2 isolated obtained from Brazil COVID-19 cases were downloaded from Global Initiative on Sharing Avian Influenza Data-EpiCoV (GISAID-EpiCoV) platform (https://www.gisaid.org/) (Shu and McCauley, 2017). Only sequences submitted up to June 30, 2021, and complete genomes (above 29,000 bp) were included. The high coverage filter was also applied to ensure acceptable quality. According to GISAID, high coverage means that only entries with less than 1% of undefined bases (NNNs) and no insertions and deletions unless verified by the submitter are tolerated. Sequences with an unidentified division were also excluded from the final dataset. The sequences were downloaded in FASTA format. The annotated reference genome sequence of the SARS-CoV-2 isolate Wuhan-Hu-1 was retrieved from the NCBI database (Accession Number: NC_045512.2).

### 2.2) Data processing

A total of 16,953 SARS-CoV-2 complete genome sequences from Brazil were obtained. To perform and for the variant calling analysis the data were grouped according to Brazilian regions (Central-West = 939; Northeast = 1,847; North = 921; South = 1,913; Southeast = 11,333). The genomes were separated by region and were aligned against the SARS-CoV-2 reference genome using Minimap2 aligner (Li et al., 2018). The SAM files from the alignments were sorted, converted to BAM and indexed using Samtools V1.9 (Li et al., 2009). The BAM file was subjected to bcftools mpileup and bcftools call (part of the samtools framework) to call variants and generate genomic VCF files. The bcftools filter was then used to filter called variations and to derive the final VCF file. The Variant Effect Predictor (VEP) was used to assess functional effects of detected variants on SARS-CoV-2 transcripts (McLaren et al., 2016).

### 2.3) Dynamics of SARS-CoV clades

Genomic surveillance of SARS-CoV-2 across Brazilian regions was performed using the Nextstrain (https://nextstrain.org/ncov), an open-source program which generates updated phylogeny with interactive visualization of publicly available SARS-CoV-2 genomes. The pipeline includes subsampling, alignment, maximum-likelihood phylodynamic analysis, temporal dating of ancestral nodes, discrete trait geographic reconstruction, and results visualization in Auspice (Hadfield et al., 2018). Because of the large number of sequenced genomes, we conduct a subsampling of Brazilian genomes from GISAID by using Nextstrain’s bioinformatics toolkit, that includes python3 scripts for preparing GISAID data for processing by Augur. This was conducted by using the --*subsample-max-sequence* option to randomly sample 300 strains from states that have a large number of sequenced genomes by using the *--query* option.

The Brazilian virus lineages were identified using Pangolin v3.1.5 as implemented on June 25th 2021 (Rambaut et al., 2020). Additionally, metadata of all SARS-CoV-2 genomes submitted on GISAID database were accessed in June 30, 2021 by using the complete genomes and high-coverage filter and the genomic clades were inferred according to its nomenclature system at the time of data collection.

### 2.4) Mathematical model to estimate the rate of genome mutants and variations between global regions

A total of 1,070,424 SARS-CoV-2 complete and high-coverage genome sequences were obtained and divided by continents: 11,574 genomes from South America; 14,986 from Oceania; 678,977 from Europe; 60,777 from Asia; 296,009 from Europe; and 8,101 from Africa. The data were processed following the conditions below and shown in **Supplementary material (Table 2 and 3)**: (i) An estimate of how many genomes are hypothetically necessary to obtain a new mutant; (ii) To compare the continents, we created an estimate index for the growing rate of variation; and (iii) To compare countries from the same regions and prove the hotspots to new variants.

#### (i) Estimates mutants *per* sequencing

we put at the center of the test the major sequencing region (Europe) comprising 1,000,285 genomes identifying a maximum of 956 lineages in 49 countries. To obtain how many genomes are necessary to obtain a new mutant, we use a factor correction to equalize the values of lineages if all the regions are hypothetically sequenced at the same rate. The factor correction is: [(ΣGE/ΣG)*(ΣLE/ΣL)], considering:

⍰ Sum of total genomes used after filtering = ΣG
⍰ Sum of total genomes from Europe = ΣGE
⍰ Sum of total lineages = ΣL
⍰ Sum of total lineages from Europe = ΣLE

Also, we apply the same logic for lineage correction and estimate the number of genomes necessary using a correction rate to identify a new variant using as a determinant variable the maximum of lineages, genomes and country.

#### (ii); (iii) Index growing variants (IGV)

⍰ Lineages per country = l
⍰ Sum of total lineages from each region: L

The index was estimated using the Shannon index variations ln(l/L) and sum products from both matrices (comparing each country, estimating the variation inside the region).

## 3) Results

### 3.1) Distribution of SARS-CoV-2 mutations in Brazil

We have evaluated the distribution of SARS-CoV-2 mutations across the five Brazilian geographical regions. In total, 42 SNVs were found across the 16,953 SARS-CoV-2 genomes sequence (**Figure 1** and **Figure 2**). 32 SNVs were found in genomes obtained from all five Brazilian regions. These mutations were found at positions C241T, T733C, C2749T, C3037T, C3828T, A5648C, A6319G, A6613G, C12778T, C13860T, C14408T, G17259T, C21614T, C21621A, C21638T, G21974T, G22132T, A22812C, G23012A, A23063T, A23403G, C23525T, C24642T, G25088T, T26149C, G28167A, C28512G, A28877T, G28878C, G28881A, G28882A, G28883C (**Figure 1** and **Figure 2**). Nine SNVs were shared across genomes from three regions (Central-West, Northeast and South) with allele frequencies ranging from 23.2% (A12964G in Central-West) to 43.5% (C12053T in Northeast). The nine mutations were found at positions C100T, T10667G, C11824T, A12964G, C12053T, C28253T, G28628T, G28975T, C29754T (**Figure 2**). Only one SNV was shared across genomes from Central-West, North and Southeast (T29834A) with allele frequency of 32.3%, 26.7% and 58.2%, respectively for each region (**Figure 2**).

**Figure 1.**
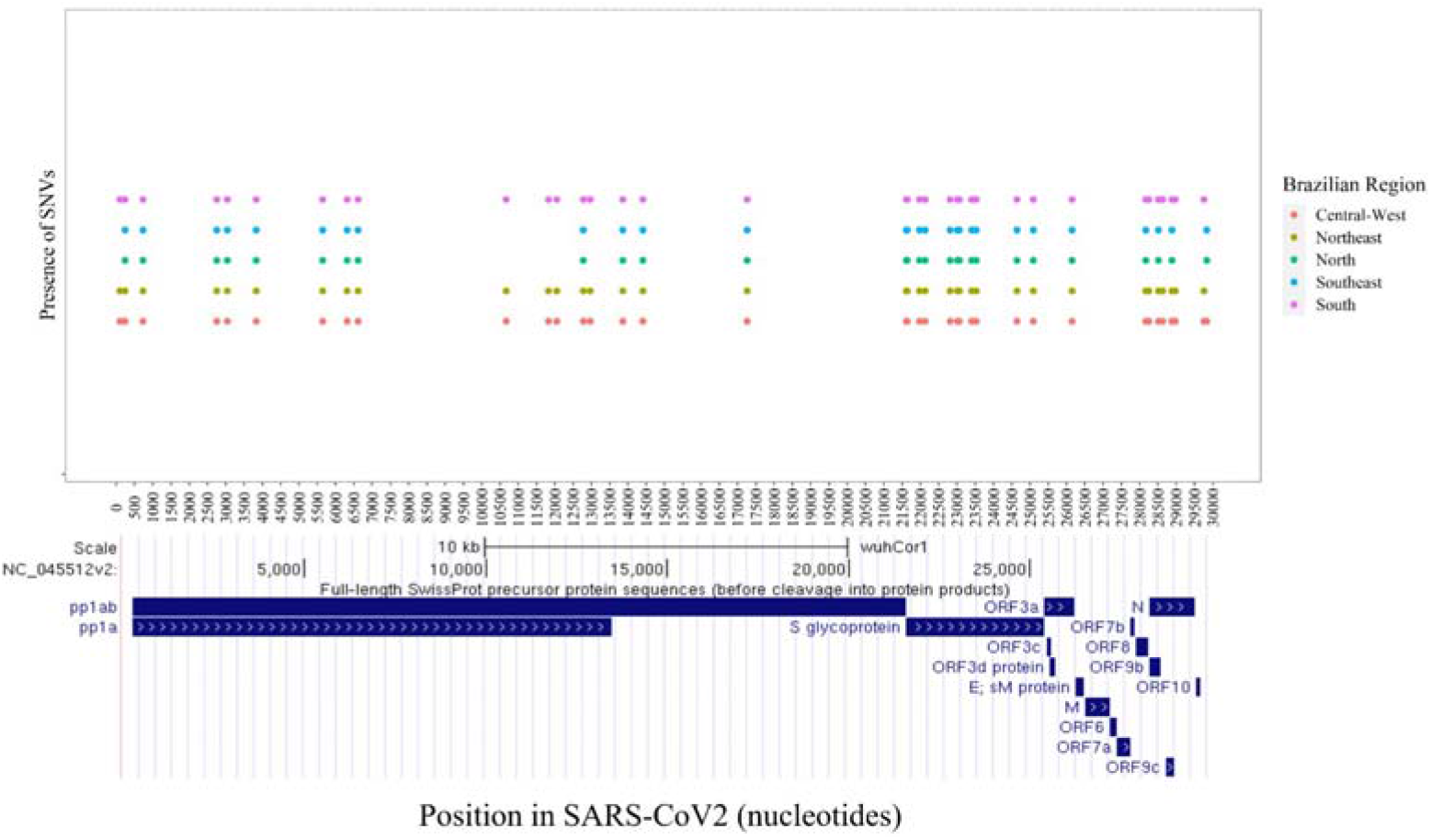
(Upper) Plot of all variants found by regions along SARS-CoV-2 nucleotide positions. (Lower) Snapshot of SARS-CoV-2 ORFs.

**Figure 2.**
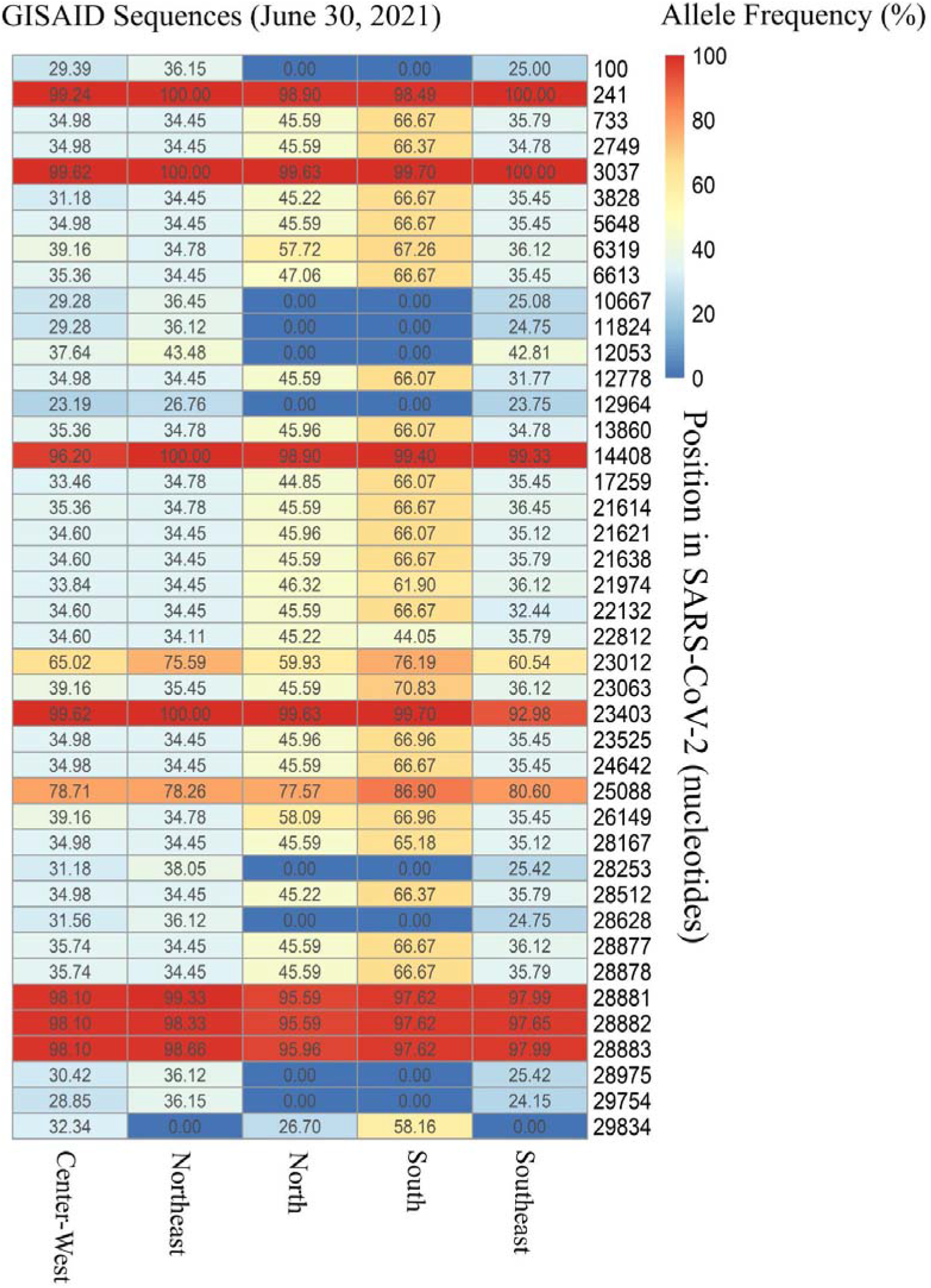
Allele frequency (plotted as percentage) of founder variants collected from Brazilian SARS-CoV-2 sequences on GISAID-EpiCoV.

Amongst the 42 found SNVs, 38 were in coding regions whereas two were in 5’UTR (C100T and C241T) and the other two in 3’UTR (C29754T and T29834A). Eleven mutations in the coding regions were synonymous or silent and 27 mutations are predicted to cause amino acid substitutions (missense variants). Among the 27 missense variants, 44.4% are located in the S gene (12-point mutations: C21614T [L18F], C21621A [T20N], C21638T [P26S], G21974T [D138Y], G22132T [R190S], A22812C [K417T], G23012A [E484N], A23063T [N501Y], A23403G [D614G], C23525T [H655Y], C24642T [T1027I], G25088T [V1176F]). It’s also important to point out that all SNVs detected in the S gene were missense mutations. Furthermore, seven missense mutations (25.9%) were located in the N gene (C28512G [P80R], G28628T [A119S], A28877T [S202C], G28878C [S202T], G28881A [R203K], G28883C [G204R], G28975T [M234I]). The ORF1ab comprises approximately 67.0% of the genome encoding 16 nonstructural proteins and had a total of six (22.6%) missense mutation (C3828T [S1188L], A5648C [K1795Q], T10667G [L3468V], C12053T [L3930F], C14408T [P4715L], G17259T [E5665D]). The ORF3a and ORF8 accounted for a total of one each; T26149C [S253P] and G28167A [E92K], respectively. In addition, nine synonymous mutations were observed in the ORF1ab (T733C, C2749T, C3037T, A6319G, C11824T, C12778T, A12964G, C13860T), and only one in the ORF8 gene (C28253T) and N gene (G28882A). Additionally, the gene E encoding envelope protein, the ORF6, ORF7a and ORF7b were conserved and did not carry any mutation (**Figure 3**).

**Figure 3.**
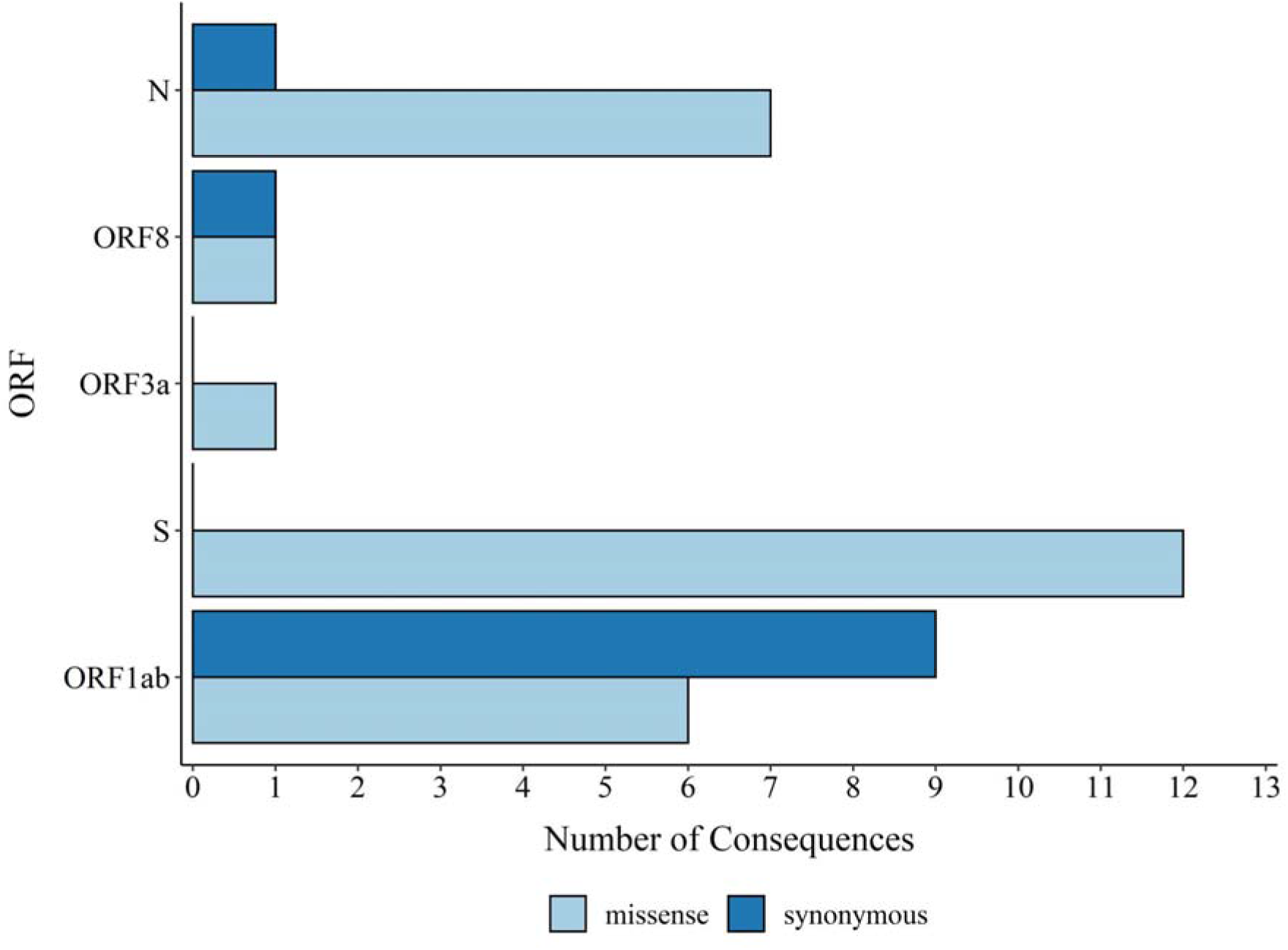
Missense and synonymous variant consequences per ORF obtained with the Variant Effect Predictor.

By reassigning sequences to the SARS-CoV-2 lineages according to the Pangolin software analysis, we observed the presence of 61 SARS-CoV-2 lineages across Brazilian regions (**Figure 4**). The majority of Brazilian genome sequences from GISAID used in this study belonged to lineages Gamma or P.1 (N=10,642; 62.8%), Zeta or P.2 (N=1,988; 11.7%), B.1.1.28 (N=1,346; 7.9%), B.1.1.33 (N=1,275; 7.5%), Alpha or B.1.1.7 (N=416; 2.5%) and P.1.2 (N=316; 1.9%), respectively. A total of 38 lineages were found in the Southeast region of Brazil, whereas 21 were found in the South. In the Northeast regions 29 lineages were found. In the North and Central-West were found nineteen and eighteen lineages, respectively.

**Figure 4.**
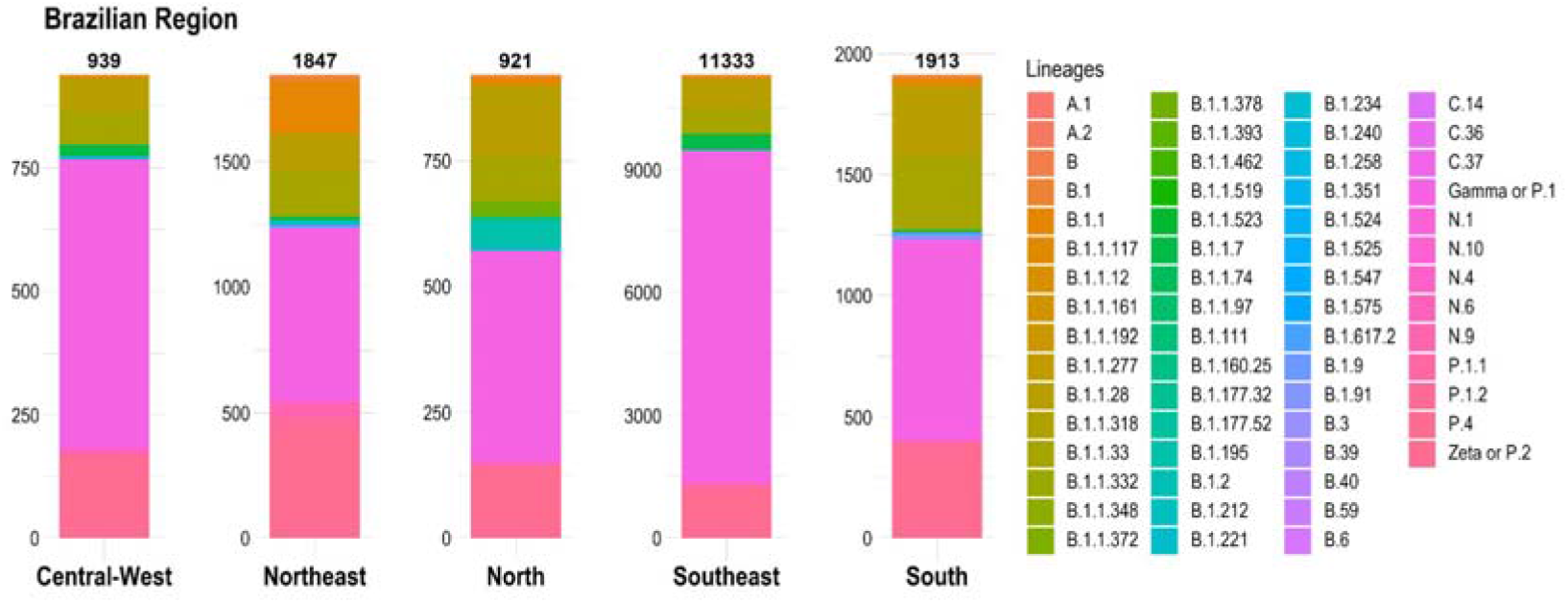
Prevalent lineages across Brazilian regions considering 16,953 genome sequences available in the GISAID database.

### 3.2) Distribution of SARS-CoV-2 clades and lineages/variants in Brazil

Regarding distribution between the five Brazilian regions, sequences from the Southeast region comprised a large proportion of Gamma (8,123 genomes), Zeta (941 genomes), B.1.1.28 (707 genomes), B.1.1.33 (644 genomes) and Alpha (374 genomes). In the South region the most prevalent lineage was Gamma (827 genomes), followed by Zeta (336 genomes), B.1.1.33 (301 genomes), B.1.1.28 (283 genomes) and P.1.2 (59 genomes). In the Northeast region, Gamma is also the most prevalent lineage (689 genomes), followed by Zeta (417 genomes), B.1.1 (206 genomes), B.1.1.33 (180 genomes) and B.1.1.28 (145 genomes). In the Central-West region the most prevalent lineages were Gamma (585 genomes), Zeta (158 genomes), B.1.1.28 (69 genomes), B.1.1.33 (63 genomes) and Alpha (23 genomes), respectively. The North region apparently has slightly different dynamics with the Gamma being the most prevalent (418 genomes), followed by B.1.1.28 (142 genomes), Zeta (136 genomes), B.1.1.33 (87 genomes) and B.1.195 (51 genomes) (**Figure 4**).

Besides, a state-by-state view shows a high distribution of the Gamma variant of concern (VOC) and the Zeta variant of interest (VOI) across almost all Brazilian states. The Gamma and Zeta lineages have been represented by genomes from twenty-six states and have not been registered only on Mato-Grosso which can be related to the extremely low sequencing rate in this state (see **Supplementary Table 1**).

To date, twenty major clades of SARS-CoV-2 were defined by Nextstrain (https://github.com/nextstrain/ncov [19A, 19B, 20A, 20B, 20C, 20D, 20E (EU1), 20F, 20G, 20H (Beta, V2), 20I (Alpha, V1), 20J (Gamma, V3), 21A (Delta), 21B (Kappa), 21C (Epsilon), 21D (Eta), 21E (Theta), 21F (Iota), 21G (Lambda) and 21H]), based on global frequency and characteristic mutational events observed in the genomes. A total of 5,351 Brazilian genomes representing all states were used for phylogenetic analysis (**Figure 5A**). The genomes were found to be classified under nine clades (19A, 19B, 20B, 20C, 20D, 20I (Alpha, V1), 20J (Gamma, V3), 21A (Delta) and 21D (Eta) and the phylogenetic examination showed that the majority belonged to clade 20B (N=2,724; 50.91%) and 20J (Gamma, V3) (2,516; 47.21%) (**Figure 5A**). As it is possible to see in **Figure 5B**, this was probably a reflection of the sequenced genomes from the beginning of the pandemic.

**Figure 5.**
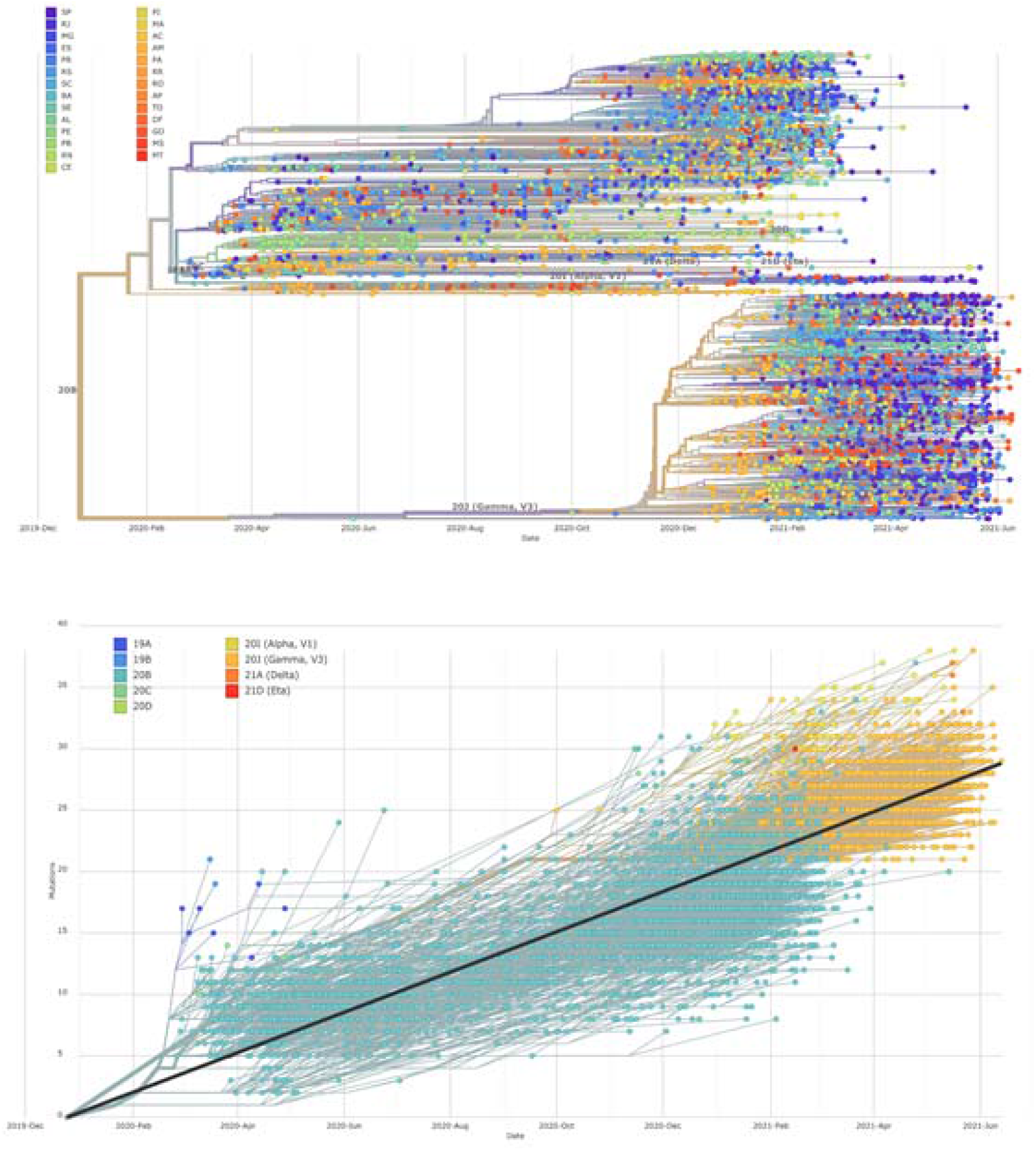
Phylogenetic tree and the clades assigned for the 5,351 brazilian SARS-CoV-2 genomes. **A**. Colored by state. **B**. Colored by clade.

It’s important to point out that different nomenclatures for SARS-CoV-2 have been proposed, including by *Nextstrain*, *cov-lineages.org* (Rambaut et al., 2020) and GISAID. Because of that we also looked at the assigned clades on the GISAID database. According to data from the GISAID database, within a year of emergence, SARS-CoV-2 has evolved into nine clades, including L, to which virus reference strains belong, S, V, G, GH, GR, GV, GRY and O (Hamed et al., 2021). The subsampled distribution of GISAID clades across Brazilian regions is shown in **Figure 6.** Overall, the clade GR (N=16,142; 95.2%) was the most prevalent among the SARS-CoV-2 genomes submitted from Brazilian regions, followed by GRY (N=375; 2.2%) and G (N=245; 1.4%). Less common clades including V, GV, S and L were identified in 0.04%, 0.02%, 0.02% and 0.02% of the submitted genomes, respectively.

**Figure 6.**
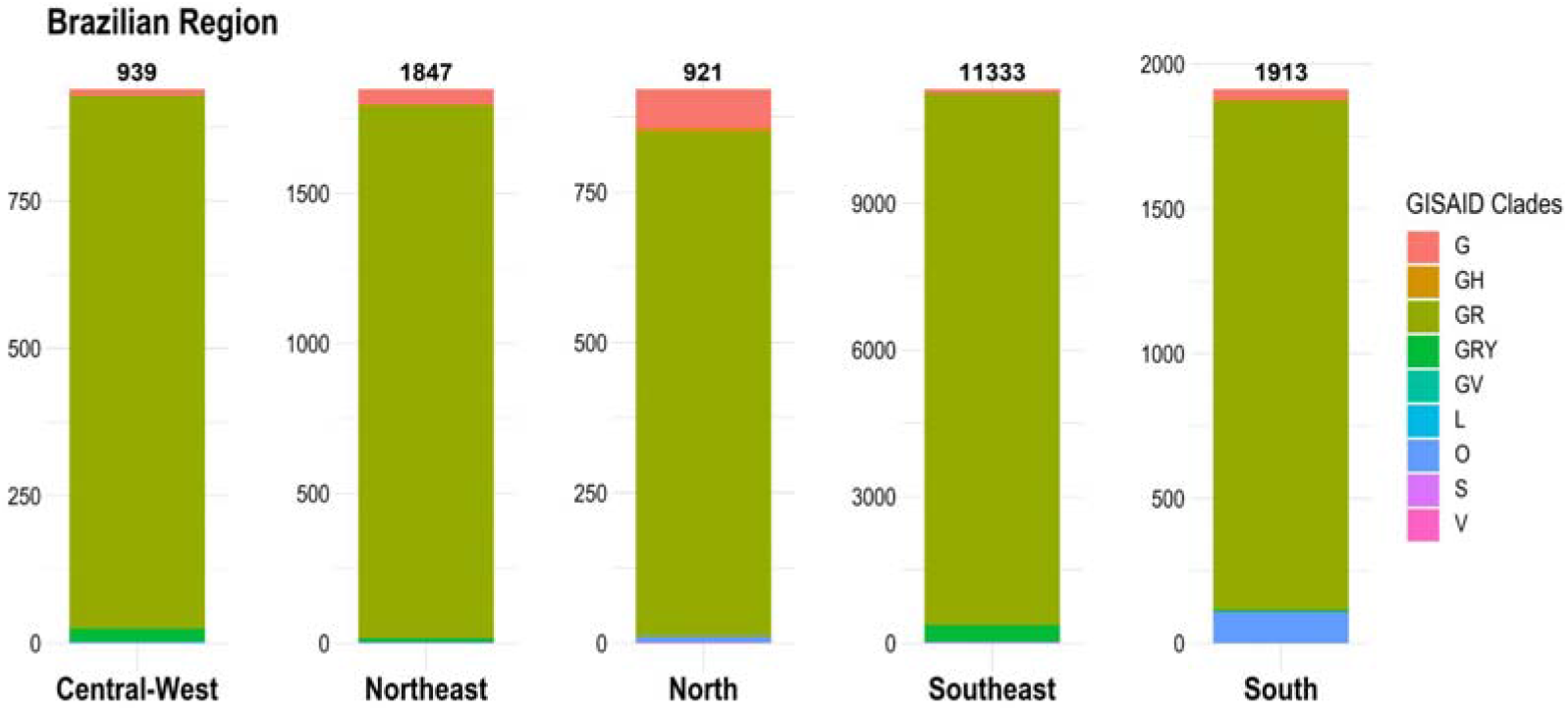
Subsampled distribution of GISAID clades across 16,953 Brazilian genome sequences.

In addition, analysis based on the chronological distribution of SARS-CoV-2 clades in Brazil showed that clade G was predominant at the beginning of the pandemic. However, this could partially be an effect of the small number of sequenced genomes. Since this initial stage the clade GR increased rapidly and stabilized 74.5% in March 2020, 89.2% in April 2020, 86.9% in May 2020 and increased further to become the most frequent clade with more than 90% in June 2020 (**Figure 7**).

**Figure 7.**
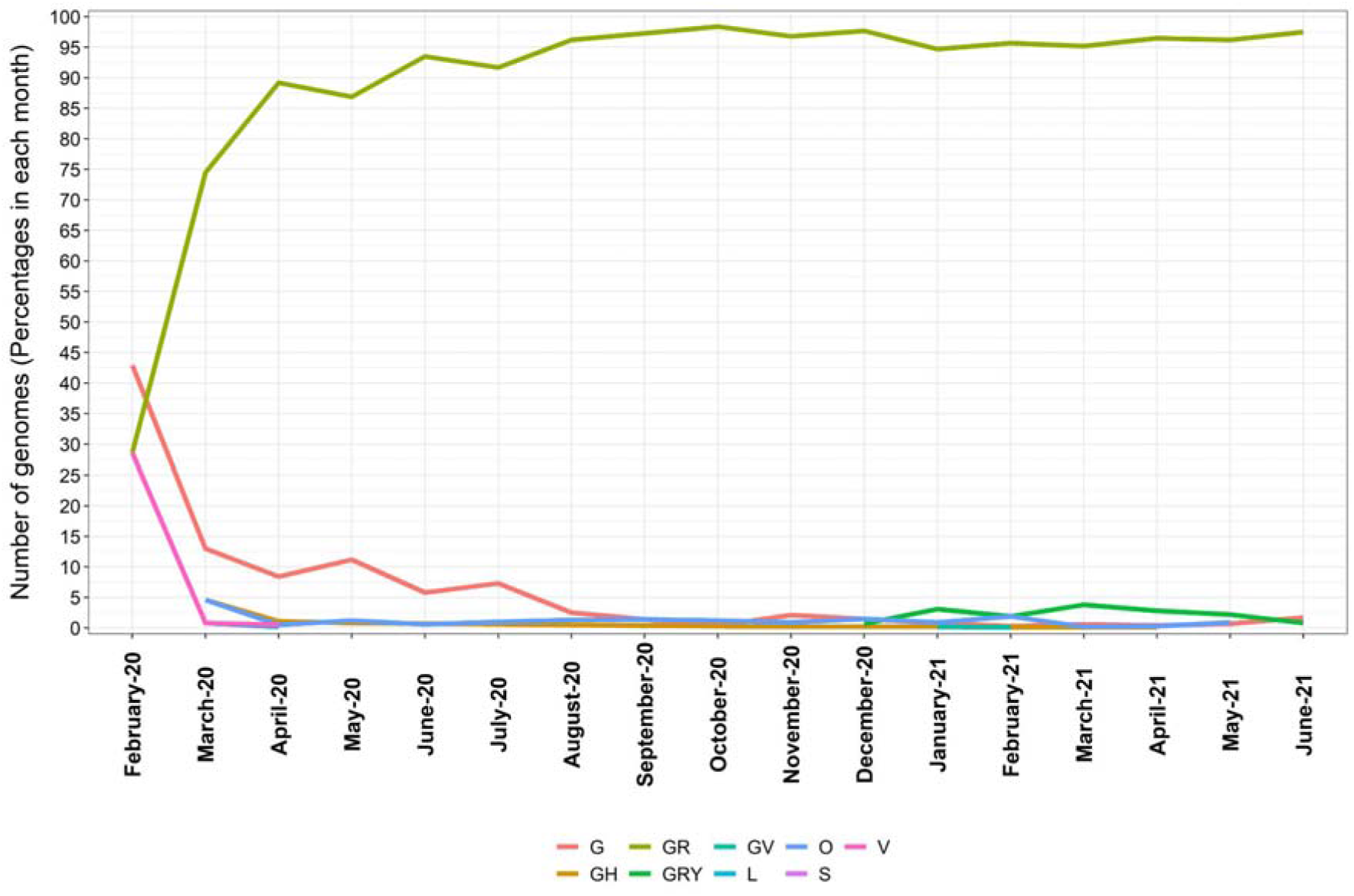
Chronological distribution of SARS-CoV-2 clades in the period from February 2020 till June 2021. Values expressed as percentages of the predominant clade in each month.

### 3.3) Numbers of genomes to identify a new variant

Regarding the published data obtained from GISAID, one question remains unclear for the authors. Is it possible to estimate how many genomes are necessary to identify a new mutant? The amount of sequences processed by countries is relative to investment and laboratory resources. On the other hand, it is possible to mathematically equalize the obtained numbers of data and create a numeric value to correct it. We downloaded 1,629,158 sequences and grouped them into: South America, North America, Europe, Africa, Asia and Oceania (**Supplementary Table 2**). Europe in total, sequenced about 1,000,285 genomes of SARS-CoV-2, identifying 956 Pango lineages in 49 countries. The differences regarding genomic surveillance and viral spread are clearly inside Europe, where the United Kingdom sequenced 41.6% of available genomes. Understanding the genome variability and evolution of genomes is fundamental to mount an effective response to contain the pandemic, however it requires governmental efforts and scientific support. We already know that these difficulties are common in other countries, leading to a gap to determine the hotspot locations by number of sequences. In **Table 1** below, we calculated how many genomes are necessary to identify a new variant, considering the quantity of genomes available. The data (**Supplementary Table 2** and **3** showed the estimated index and applied formulas) were equalized with a correction value estimated as 0,06485746617. The column G/L shows the data previously equalized. This analysis focuses on relevance to regions as hotspots for new variants, in cases of the number G/L is lower than others, indicating the major probability to find a new mutant each the number reported. On the other side, the last column (**Table 1**) showed a diversity analysis between countries of each region, considering the maximum number of lineages of each country between geographical regions. This index represented by alpha diversity (**Table 1**) shows the high diversity of lineages across Europe, numerically represented as 8.4996. In the range of each country, Europe varies from 0.0072 to the highest value 0.3676 represented by France. In the second range of diversity, Africa reported 6.4225 in which South Africa presented the highest rate (0.3381) followed by Asia 5.9559. Belonging to Asia, the model put India 0.3677 in the hotspot region to new variants followed by Japan (0.3542). Across South America, the index was 3.4064, probably explained by the low availability of data and less proportion of cases in some countries belonging to this geographic area. Analyzing each country in these regions, we observed that Chile (0.3679) and Brazil (0.3650) represented, according to genomes available, the same and the highest tax of variability expressed in parenthesis. North America (1.6557) demonstrates low levels of diversity, in which Canada is the hotspot (0.3574). Oceania, as expected, represented a minimum tax of variation estimated, showing an index of 0.9893.

**Table 1.** Total number of genomes after filtering quality deposited and downloaded from Gisaid. The data showed predictivity values to identify one new variant as a function of sequences per region.

| **Region** | **n of Genomes** | **n max of Lineages** | **G/L \*** | **n of countries** | **Diversity region index** |
| --- | --- | --- | --- | --- | --- |
| South America | 26,257 | 189 | 25.28 | 15 | 3.6906 |
| Oceania | 15,484 | 249 | 24.85 | 6 | 0.9893 |
| Europe | 1,000,285 | 956 | 293.21 | 49 | 8.4996 |
| Asia | 112,901 | 512 | 49.01 | 38 | 5.9559 |
| North America | 477,842 | 732 | 188.30 | 17 | 1.6557 |
| Africa | 11,873 | 239 | 13.99 | 38 | 6.4225 |

\*G/L after applied correction factor

## 4) Discussion

As SARS-CoV-2 continues to circulate in the human population after more than one year of pandemic, it is natural to observe genetic differences between SARS-CoV-2 strains sampled in various locations. This is the largest study focused on genome-wide mutational spectra covering nucleotides, amino acids and deletion mutations in 16,953 complete and high coverage SARS-CoV-2 genomes from the five Brazilian geographical regions. Furthermore, this preliminary and crucial analysis of the Brazilian SARS-CoV-2 genome shows the increase in the number of mutations.

Viral mutations are probabilistic events through the random transmission between infected people. The viral load is variable and depends on such factors as the course of infection and host immunity. Some individuals are “super spreaders” which means that the behavioral and environmental variables are relevant to infectivity, increasing the successful transmission (Cave, 2020). As of date, we have around 19 million cases. So, here we are analyzing only ~0.05% of reported cases, comprising a snapshot of SARS-CoV-2 mutations status in Brazil.

A number of previous studies have examined variants within SARS-CoV-2 isolates. Rouchka et al. (2020) reported clustered groups of sequences showing geographical similarities, suggesting clusters of similar transmission in both time and viral strains. Resende et al. (2021) showed that B.1.1.28 (E484K) is present in several states from the South, Northeast, and North Brazilian regions and dates its origin to August 27th, 2020 (July 14th - September 18th). These findings documented a classical SARS-CoV-2 reinfection case with the emerging Brazilian lineage B.1.1.28 (E484K). Additionally, provide evidence of this emerging Brazilian clade’s geographic dissemination outside the Rio de Janeiro state. Naveca et al. (2021) reported a preliminary genomic analysis of SARS-CoV-2 B.1.1.28 lineage circulating in the Brazilian Amazon region and their evolutionary relationship with emerging and potential SARS-CoV-2 Brazilian variants harboring mutations in the RBD of spike protein. Phylogenetic analysis of 69 B.1.1.28 sequences isolated in the Amazonas state revealed the existence of two major clades that have evolved locally without unusual mutations in the spike protein from April to November 2020. In Africa, Motayo et al. (2020) showed the high prevalence of the D614 spike mutation in order to 82% between sequences analyzed.

More than identifying the mutations, these analyses allow continuous research focused on mapping how amino-acids changes affect antibody binding. In a recent study, Nonaka et al. (2021) report the first case of reinfection from genetically distinct SARS-CoV-2 lineage presenting the E484K spike mutation in Brazil, a variant associated with escape from neutralizing antibodies. The mutations on the RBD domain enhanced ACE2 binding, promoting viral infectivity and maybe disrupting neutralizing antibodies (NAb) binding to evade the host immune response. Antibodies targeting RBD have been used and developed as therapeutics and are known as the major contributors to NAb responses. To control viral infection a robust humoral immune response is essential in populations. Scanning mutations is important to map the RBD changes and used to predict escape mutations in antibody epitopes (Hou et al., 2020).

Nevertheless, SARS-CoV-2 genomes sequenced in Brazil until now were clustered into at least nine major clades, as defined by the GISAID database. Of them, Clade GR was the most frequently identified in Brazilian genomes, followed by GRY and G. Since Clade G (Lineage B.1), defined by the spike protein’s D614G mutation, was identified, it rapidly predominated in many locales where it was found. Theoretical evidence suggests that mutations in the viral spike may be linked to altered potential for host cell membrane fusion, which should result in increased person-to-person transmission and pathogenicity (Brufsky et al., 2020; Korber et al., 2020; Walls et al., 2020). Sub cluster of clade G then started to split into GR, GH, GV and more recently into GRY. Similar to what was observed recently by Hamed et al. (2021), the analysis of the distribution of SARS-CoV-2 genomes across continents showed that there was much expansion in the number of sequence genomes that were clustered into the GR and GRY clade compared to clade G, suggesting higher fitness for transmission by the newer clades compared to their ancestral one.

Here we evaluated the distribution of SARS-Cov-2 mutations across the five Brazilian geographical regions, showing different allelic frequencies with similar general distribution of all variants across different regions. We also showed the presence of 27 missense variants in the entire genome, the majority (44.4%) being present in the spike gene. Based on Pangolin software, we show the presence of 61 SARS-CoV-2 lineages across Brazilian regions, with a high predominance of the Gamma variant. Based on Nextstrain clades, Brazilian genomes were classified into 9 clades, with the majority belonging to clade 20B (N=2,724; 50.91%) and 20J (Gamma, V3) (2,516; 47.21%). In GISAID clades, there are also 9 clades, with a predominance of the GR clade (95.2%).

Finally, we estimate the number of genomes needed to generate a variant and what is the ratio of this index by continent. The aim of diversity analysis was to show the hotspot regions in a printed scenario of pandemics. While we have a high genomic diversity in Europe given the large number of sequenced genomes, Africa is emerging as a hotspot for new variants. Asia is the third continent in terms of diversity. In South America, Brazil and Chile, they have rates similar to South Africa and India. These numbers concern the attention of hotspot regions to emerging new variants and suggest rapid vaccination protocols to control the spread of virus. All of these data taken together demonstrate a wide circulation of SARS-CoV-2 throughout the continents, a favorable scenario for viral multiplication. The genomic surveillance showed a potential tool to monitor the circulation of SARS-CoV-2 and understand the biological characteristics of the viral genome.

## Acknowledgements

We offer our deepest thanks to the Global Initiative on Sharing Avian Influenza Data-EpiCoV (GISAID-EpiCoV) for the data made available and updated on the platform (https://www.gisaid.org/). A full list acknowledging the authors publishing data used in this study can be found in the following file: **Supplementary Table 4** (844 Kb). U.J.B.S. and R.N.S. are granted a post-doctoral scholarship (DTI-A) from CNPq. This work was supported by Rede Corona-ômica BR MCTI/FINEP (http://www.corona-omica.br-mcti.lncc.br) affiliated to RedeVírus/MCTI (FINEP = 01.20.0029.000462/20, CNPq = 404096/2020-4). The study was also supported by FAPERGS (21/2551-0000081-3). We would like to thank Anderson Brito for his help with the Nextstrain tool.

## Research ethics and consent

This study does not require the use of an ethics committee as we use the genomes deposited in the GISAID database.

